# Effects of Dopamine Receptor Antagonists and Radiation on Mouse Neural Stem/Progenitor Cells

**DOI:** 10.1101/2023.01.18.524632

**Authors:** Ling He, Kruttika Bhat, Angeliki Ioannidis, Frank Pajonk

## Abstract

**Background:** Dopamine receptor antagonists are psychotropic drugs that have been originally developed against psychiatric disorders. We recently identified dopamine receptor antagonists as potential anti-cancer agents and some have entered clinical trials against glioblastoma. Radiotherapy is known to cause cognitive impairment in patients receiving cranial irradiation through the elimination of neural stem/progenitor cells and subsequent loss of neurogenesis.

**Methods:** Using transgenic mice that report the presence of neural stem/progenitor cells through Nestin promoter-driven expression of enhanced green fluorescent protein, the effects of dopamine receptor antagonists alone or in combination with radiation on murine neural stem/progenitor cells were assessed in sphere-formation assays, flow cytometry and immunofluorescence *in vitro* and *in vivo*.

**Results:** We report that several dopamine receptor antagonists show sex-dependent effects on neural stem/progenitor cells both *in vitro* and *in vivo*. Hydroxyzine, trifluoperazine, amisulpride, nemonapride or quetiapine alone or in combination with radiation significantly increased the number of neural stem/progenitor cells in female neurospheres but not in male mice. Dopamine receptor antagonists either protected neural stem/progenitor cells from radiation or expanded the stem cell pool, thus indicating that this combination therapy against glioblastoma will not increase radiation-induced cognitive decline through increasing elimination of neural stem/progenitor cells and subsequent loss of neurogenesis.

**Conclusions:** We conclude that a therapeutic window for dopamine receptor antagonists in combination with radiation potentially exist, making it a novel combination therapy against glioblastoma. Normal tissue toxicity of this combination potentially differs depending on age and sex and should be taken into consideration when designing clinical trials.

**Key Points:**

- Neural stem/progenitor cells show sex-dependent sensitivity to dopamine receptor antagonists
- Dopamine receptor antagonists active against GBM increase Neural stem/progenitor cells counts

**Importance of the Study:** Combination therapy of dopamine receptor antagonists with radiation have entered clinical trials against glioblastoma but the normal tissue toxicity of this combination has not been fully explored yet. Here we present evidence that some dopamine receptor antagonists show sex-dependent effects on neural stem/progenitor cells either by protecting neural stem/progenitor cells from radiation or inducing an expansion of the stem cell pool, suggesting that this combination therapy against glioblastoma will not increase radiation-induced cognitive decline through increasing elimination of neural stem/progenitor cells and subsequent loss of neurogenesis. Normal tissue toxicity of this combination potentially differs depending on age and sex and should be further explored in clinical trials.

## Introduction

Neurotransmitters and their receptors were originally described in neuronal tissues (1, 2) but have since been found to be expressed in many solid cancers including brain tumors (3, 4). Recent studies have identified dopamine receptors as potential targets for cancer therapy and dopamine receptor antagonists (DRAs) have entered clinical trials against glioblastoma (5–7). DRAs are psychotropic drugs originally developed against a broad range of psychiatric disorders. These drugs have well described side effect profiles in normal tissues and their pharmacokinetic and pharmacodynamic properties are well established. Given their ability to cross the bloodbrain barrier, repurposing of these FDA-approved drugs for use in intracranial malignancies highlights DRAs as attractive candidates in the context of glioblastoma.

DRA treatment by itself has shown limited efficacy against some cancers but we recently reported significantly improved survival in mouse models of glioblastoma when DRAs were combined with radiotherapy (6, 8, 9). Radiotherapy is known for causing cognitive impairment in patients receiving cranial irradiation. This side effect is most pronounced in children (10) but also observed in adult patients and at least in part thought to be caused by the loss of neurogenesis (11, 12). Data on the combined effects of radiation and DRAs on neural stem/progenitor cells is not available in the literature. We recently reported that the combined treatment with radiation and DRAs eliminated glioma-initiating cells both *in vitro* and *in vivo* (6, 8, 9). In light of some similarities between neural stem/progenitor cells (NSPCs) and glioma-initiating cells (GICs) we sought to investigate whether DRAs alone or in combination with radiation adversely affect NSPCs.

## Materials and Methods

### Animals

Nestin-EGFP mice were a kind gift from Dr. Grigori Enikolopov, Cold Spring Harbor Laboratory (13). C3Hf/Sed/Kam mice were originally obtained from the MD Anderson Cancer Center. Mice were re-derived, bred and maintained in a pathogen-free environment in the American Association of Laboratory Animal Care-accredited Animal Facilities of the Department of Radiation Oncology, University of California (Los Angeles, CA) in accordance with all local and national guidelines for the care of animals. Sex of newborn animals was determined as described in (14).

### Neural stem cell culture

For neural stem cell cultures, the entire brains of the newborn Nestin-EGFP mice were harvested, minced using a scalpel and transferred to a fresh Eppendorf tube with 1 ml of 0.05 % trypsin-EDTA (Thermo Fisher Scientific, # 25300-054), mixed and incubated at 37 °C for 7 minutes. Trypsin was inhibited by adding 1 ml of trypsin inhibitor (Sigma-Aldrich, Cat# T6522). The tissue was centrifuged at 110 *x g* for 5 minutes and the supernatant was discarded. The tissue was resuspended in 200 μl of NeuroCult (Stem Cell Technologies, # 05702). The solution was mixed well to break down the tissue, diluted with 10 ml NeuroCult media and passed through a 70 μm cell strainer. The cell suspension was centrifuged at 110 *x g* for 5 minutes, the supernatant discarded, and the cell pellet resuspended in 2 ml of ACK lysing buffer (Lonza, Cat# 10-548E). After centrifugation, cells from the final pellet were cultured in 1X complete NeuroCult media (45 ml of base NeuroCult media, 5 ml of supplement media, 1 μl of EGF, 20 μl of bFGF, 3130 U/ml of heparin) under standard conditions (37 °C, 5 % CO2) and labeled as Passage #1. All experiments were performed with Passage #2 cells.

### Irradiation

Neurosphere cultures were irradiated with 0, 2 or 4 Gy at room temperature using an experimental X-ray irradiator (Gulmay Medical Inc. Atlanta, GA) at a dose rate of 5.519 Gy/min. Control samples were sham-irradiated. The X-ray beam was operated at 300 kV and hardened using a 4 mm Be, a 3 mm Al, and a 1.5 mm Cu filter and calibrated using NIST-traceable dosimetry. Corresponding controls were sham irradiated.

### Drug treatment

For *in vitro* studies DRAs were solubilized in DMSO or PBS. The neurosphere cultures were pre-treated either with 1 μM DRAs or solvent control before irradiation with 0, 2 or 4 Gy. In *in vivo* experiments mice received 5 daily i.p. injections of 20 mg/kg of trifluoperazine, 30 mg/kg of quetiapine, 20 mg/kg of hydroxyzine dihydrochloride, 1 mg/kg of nemonapride or 1 mg/kg of amisulpride.

### Flow cytometry

Passage #2 neurospheres established from the brains of Nestin-EGFP mice were harvested and dissociated into single cell suspension as described above. Single cell suspensions were either subjected to FACS (Flow Cytometry Core, Broad Stem Cell Research Center at UCLA, BD FACS ARIA) for EGFP-high, -medium, and -low cell populations or analyzed for total EGFP expression using a MACSQuant Analyzer (Miltenyi Biosciences, Auburn, CA) and the FlowJo software package (v10, FlowJo, Ashland, OR). Drug-treated samples were normalized against the corresponding solvent-treated control sample (PBS or DMSO).

### Neurosphere-formation assay

To assess self-renewal capacity, Passage #2 neurospheres from Nestin-EGFP mice were dissociated and plated in an *in-vitro* limiting dilution assay into 96-well non-treated plates containing 1X complete NeuroCult media, at a range of 1 to 1,000 cells/well. Growth factors, EGF and bFGF, were added every 3 days, and the cells were allowed to form neurospheres for 14 days. The number of spheres formed per well was then counted and expressed as a percentage of the initial number of cells plated.

### Brain dissection

Five days after drug treatment, the brains of the mice were harvested and placed on the Acrylic Mouse Brain Slicer Matrix (Zivic Instruments, Cat# BSMAA001-1). 1 mm thick coronal sections starting from 2 mm anterior to 2 mm posterior of the bregma were cut. The brain tissue was minced into very small pieces using a scalpel and the cells were isolated in Neural Stem Cell culture media as described above. Cells were then used for flow cytometric analysis to assess the percentage of Nestin-EGFP^+^ cells in different treatment groups.

### Immunofluorescence

Brains were explanted, fixed in formalin for 24 hours and embedded in paraffin. 4 μm blank sections were baked in an oven at 65 °C for 30 minutes, dewaxed in two successive Xylene baths for 5 minutes each and then hydrated for 5 minutes each using an alcohol gradient (ethanol 100 %, 90 %, 70 %, 50 %, 25 %). Antigen retrieval was performed using Heat Induced Epitope Retrieval in citrate buffer (10 mM sodium citrate, 0.05 % tween20, pH=6) with heating to 95 °C in a steamer for 10 minutes. After cooling down, the slides were blocked with 10 % goat serum containing 1 % BSA at room temperature for 30 minutes and then incubated with the primary antibody against EGFP (Abcam, ab184601, 1:100) overnight at 4 °C. The next day, the secondary antibody Alexa Fluor 488 Goat Anti-mouse IgG (H/L) antibody [1:1,000 (Invitrogen)] was applied for 60 minutes. Slides were mounted with ProLong^®^ Gold Antifade Reagent with DAPI (Cell signaling, Cat #8961S). Images were captured and merged at 20X using a digital microscope (BZ-9000, Keyence, Itasca, IL) and the percentages of Nestin-EGFP^+^ cells were quantified using Image J.

### Statistics

Unless stated otherwise all data shown are represented as mean ± standard error mean (SEM) of at least 3 biologically independent experiments. A *p*-value of ≤0.05 in an unpaired two-sided *t*-test or two-sided ANOVA test for multiple comparisons indicated a statistically significant difference.

## Results

### Dopamine receptor antagonists show sex-dependent effects on neural stem/progenitor cells in vitro

In a first set of experiments, we tested if DRAs affect the number of NSPCs *in vitro*. Nestin-EGFP^+^ NSPCs were harvested from both male and female newborn pups and cultured as neurospheres. We have previously reported that Nestin promoter driven EGFP expression in this mouse strain is correlated with self-renewal capacity (15). Passage #2 neurospheres were pretreated with DRAs at 1 μM concentrations, irradiated with 0, 2 or 4 Gy and cultured for 2 weeks during which DRAs were added 3x/week **(Figure 1A)**.

**Figure 1.**
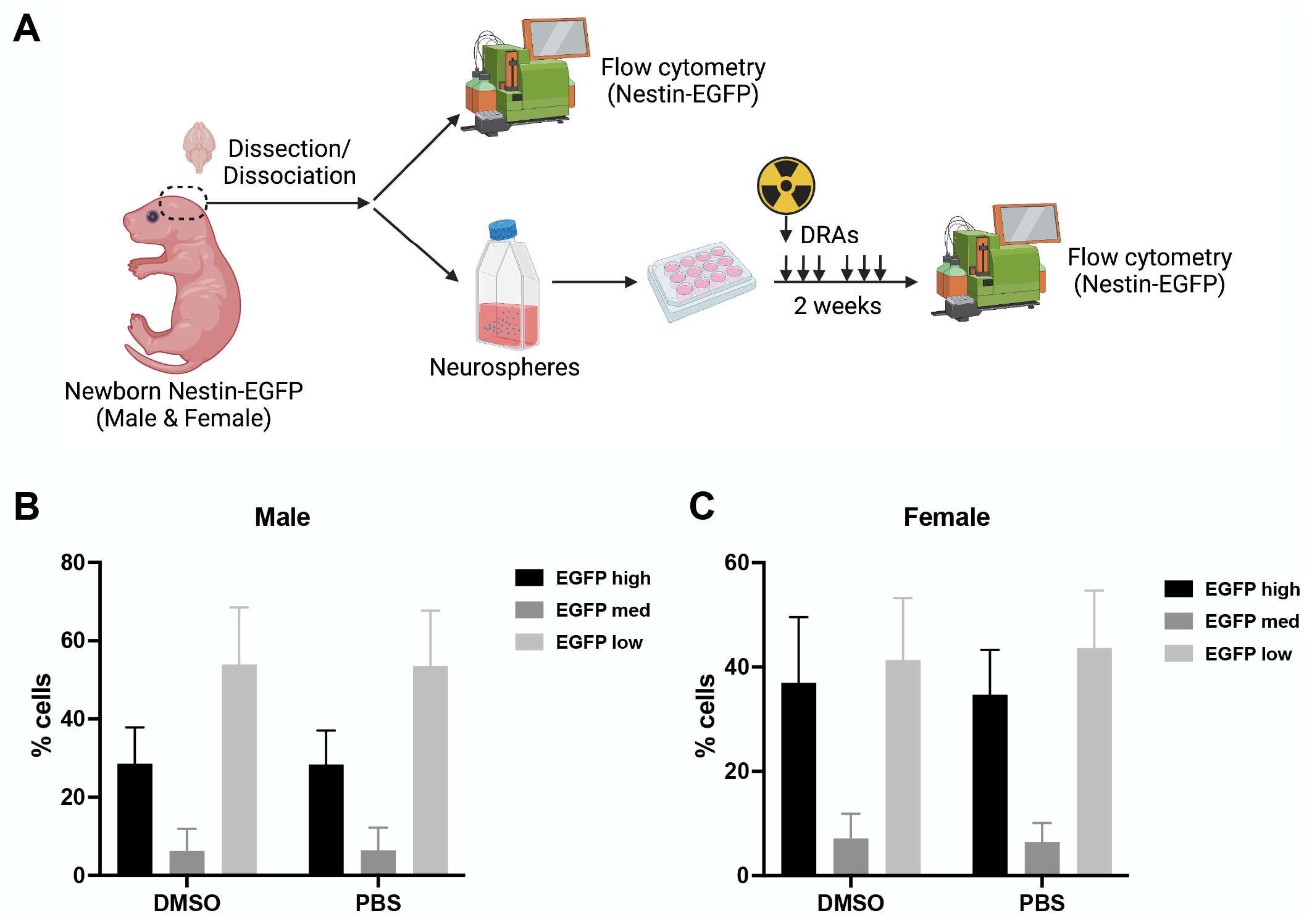
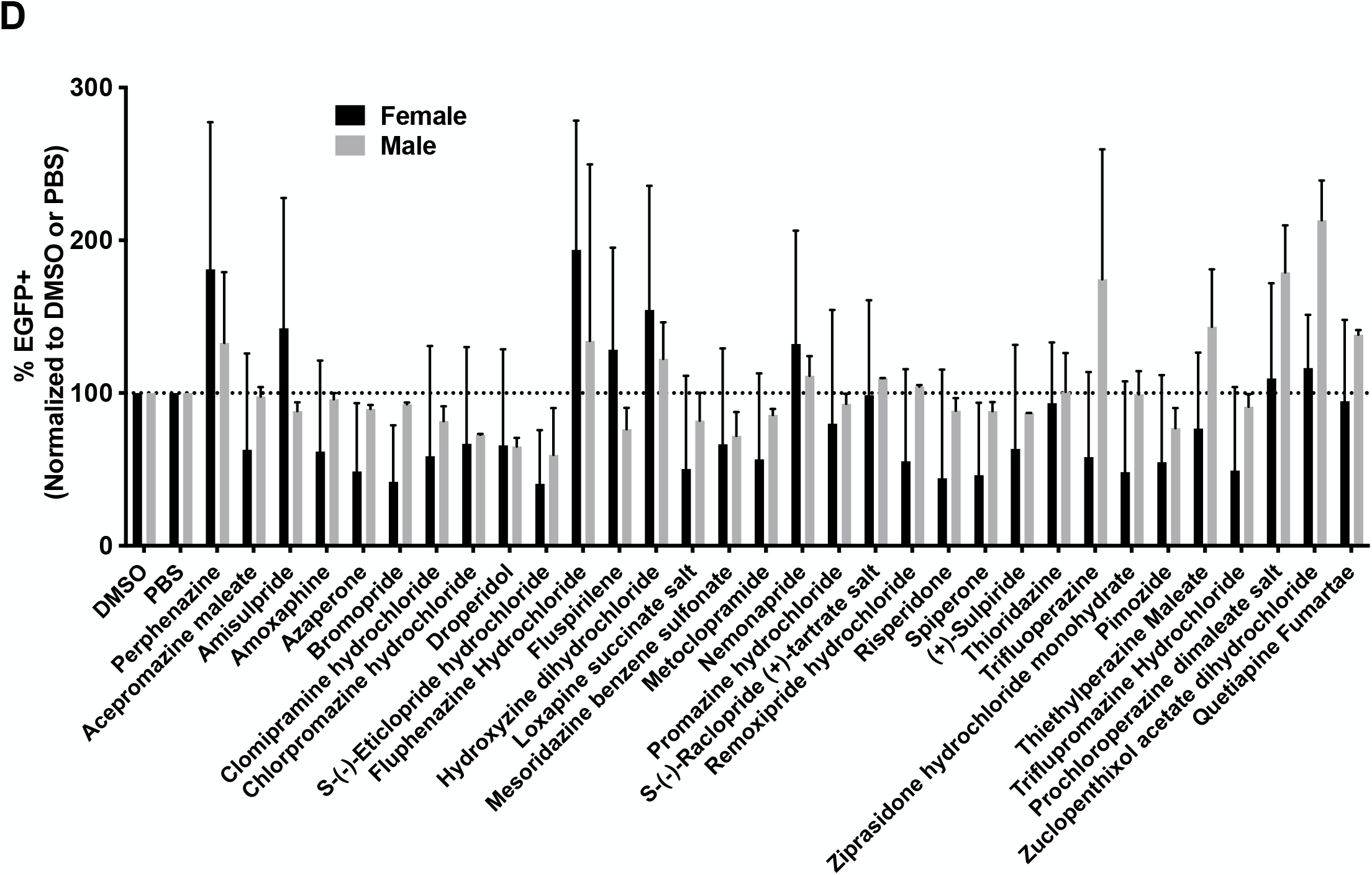
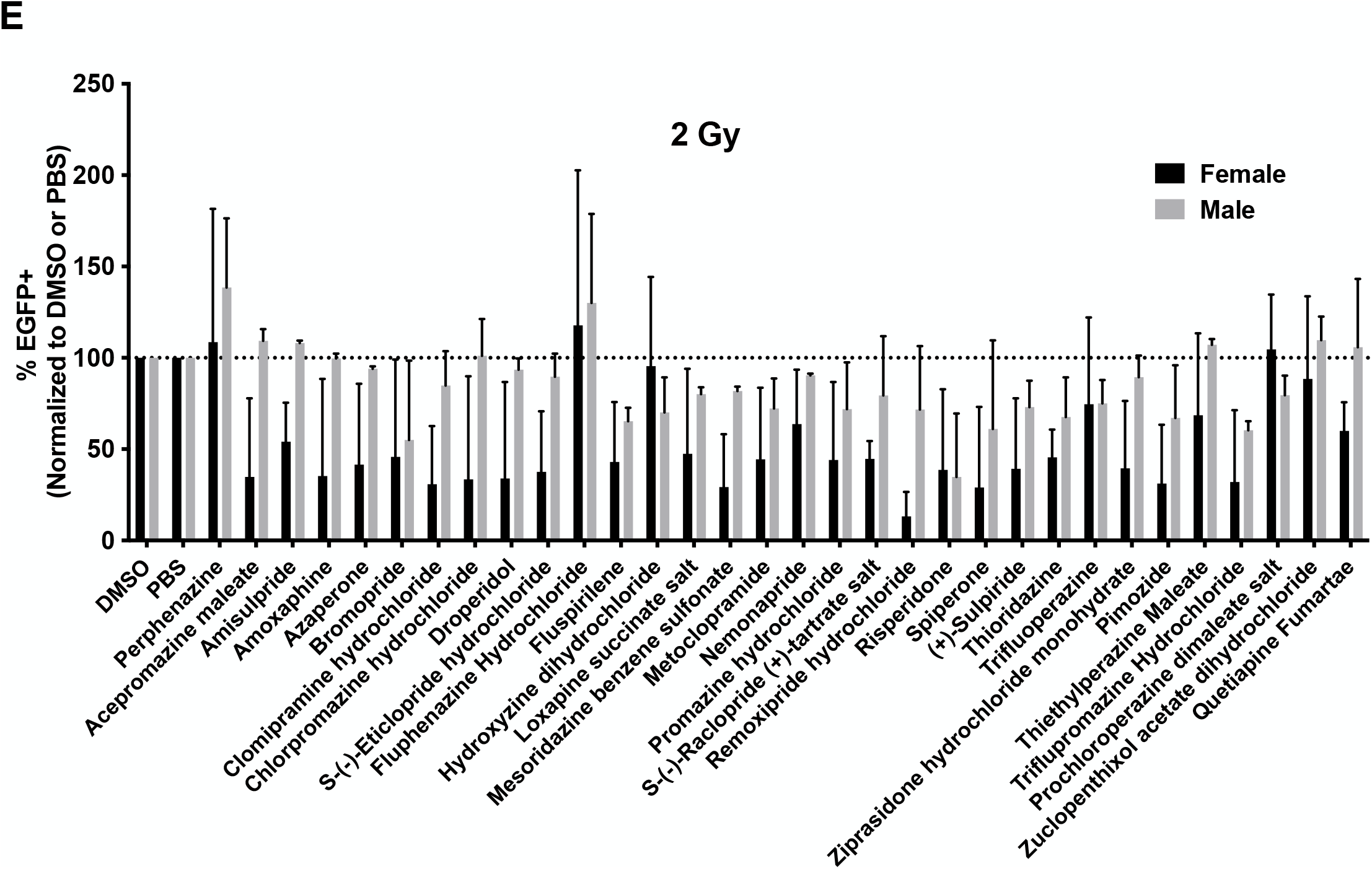
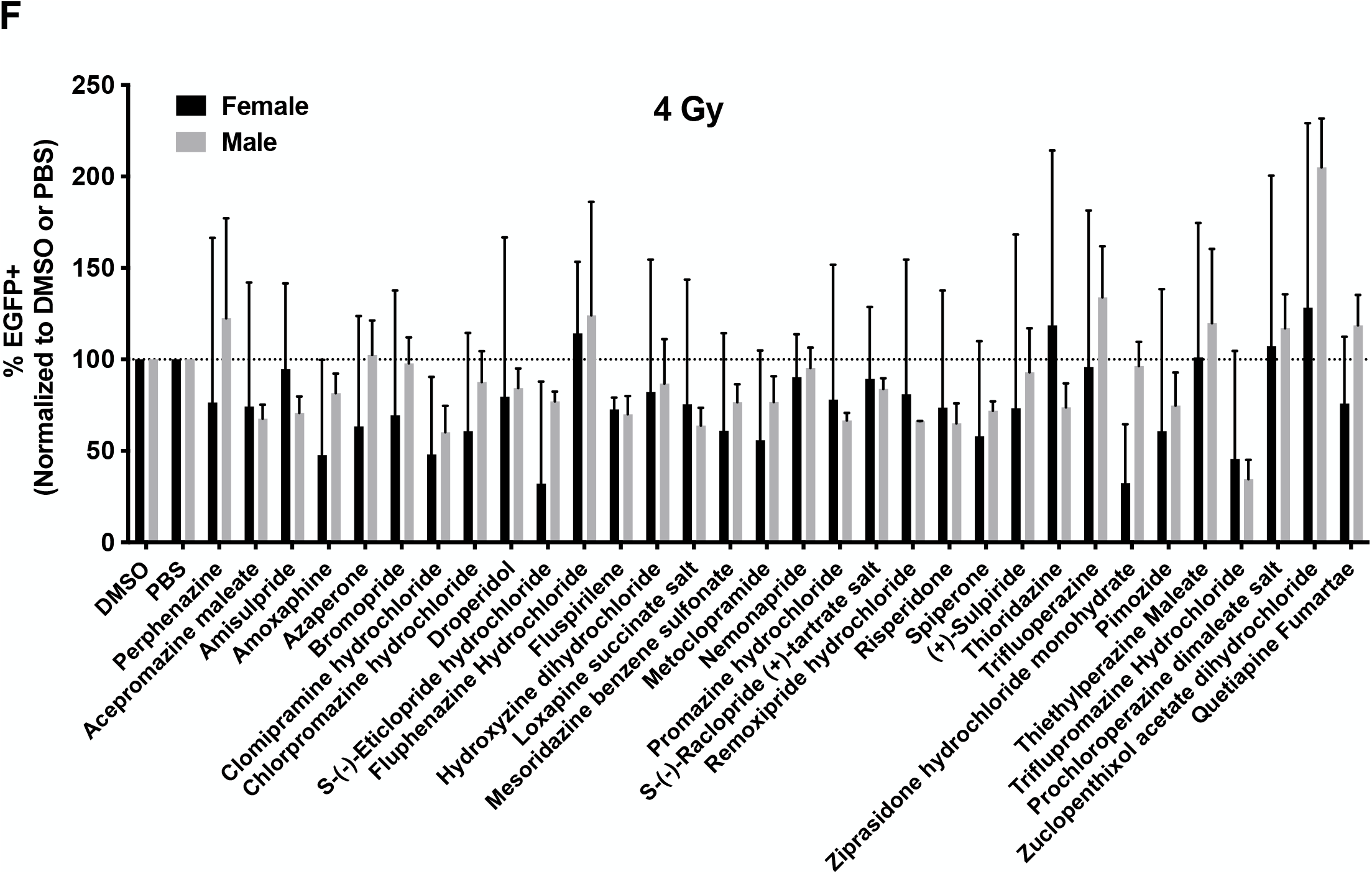
**(A)** Schematic of the experimental design underlying Figure 1. **(B)** The brains from male or female newborn pups were harvested, dissociated, and analyzed for EGFP-^high^, -^medium^ and -^low^ subpopulations by flow cytometry (N=3). **(C-E)** Neurospheres (Passage #2) from male or female newborn pups were pre-treated with either controls (DMSO/PBS) or DRAs (1 μM) and subjected to irradiation with 0 **(C)**, 2 **(D)** or 4 Gy **(E)**. DRAs were added 3 times per week for two weeks, time after which the neurospheres were subjected for flow cytometry and analyzed for the total percentage of EGFP+ cells in each sample (N=3).

At the conclusion of this experiment 2 weeks post treatment initiation we observed baseline differences in the number of Nestin-EGFP^high^ NSPCs in neurospheres established from male and female mice with female mice showing a higher stem cell content [female: 37 ± 12.6 % (DMSO), 34.7 ± 8.6 % (PBS); male: 28.6 ± 9.3 % (DMSO), 28.4 ± 8.7 % (PBS); N=3] **(Figure 1B)**. In response to DRA treatment alone we found unexpected differences between neurospheres established from male and female mice, with those from female mice in general showing larger reductions in the number of EGFP^+^ NSPCs. However, the number of EGFP^+^ NSPCs in Passage #2 neurospheres from individual animals of the same sex varied and none of the observed differential effects of DRAs between sexes reached the level of statistical significance **(Figure 1C)**. In agreement with our previous report (15), radiation predominately eliminated the Nestin-EGFP^high^ population of cells from neurospheres, consistent with the known exquisite radiation sensitivity of neural stem cells. The differences between sexes in the total number of EGFP^+^ cells in response to individual DRAs became smaller with increasing radiation doses in alignment with the generally non-sex-specific cell killing effects of ionizing radiation **(Figure 1D/E)**.

Next, we tested if the DRAs that showed the largest effects on the number of Nestin-EGFP^+^ cells in neurospheres established from either male or female mice also affected the self-renewal capacity of NSPCs from the same Passage #2 neurospheres. Cells were plated at clonal density in an *in vitro* limiting dilution assay, irradiated and treated with DRAs. The number of clonal spheres formed was counted two weeks later **(Figure 2A)**. Cells from neurospheres established from male mice showed a significant reduction in sphere-formation after irradiation with 0, 2, or 4 Gy compared to the corresponding irradiated, solvent-treated controls when treated with perphenazine, fluphenazine, thiethylperazine, prochloroperazine, or zuclopenthixol but not after treatment with hydroxyzine, trifluoperazine, amisulpride, or nemonapride. At the same time, treatment with quetiapine significantly increased sphere-formation after irradiation with 2 Gy **(Figure 2B)**.

**Figure 2.**
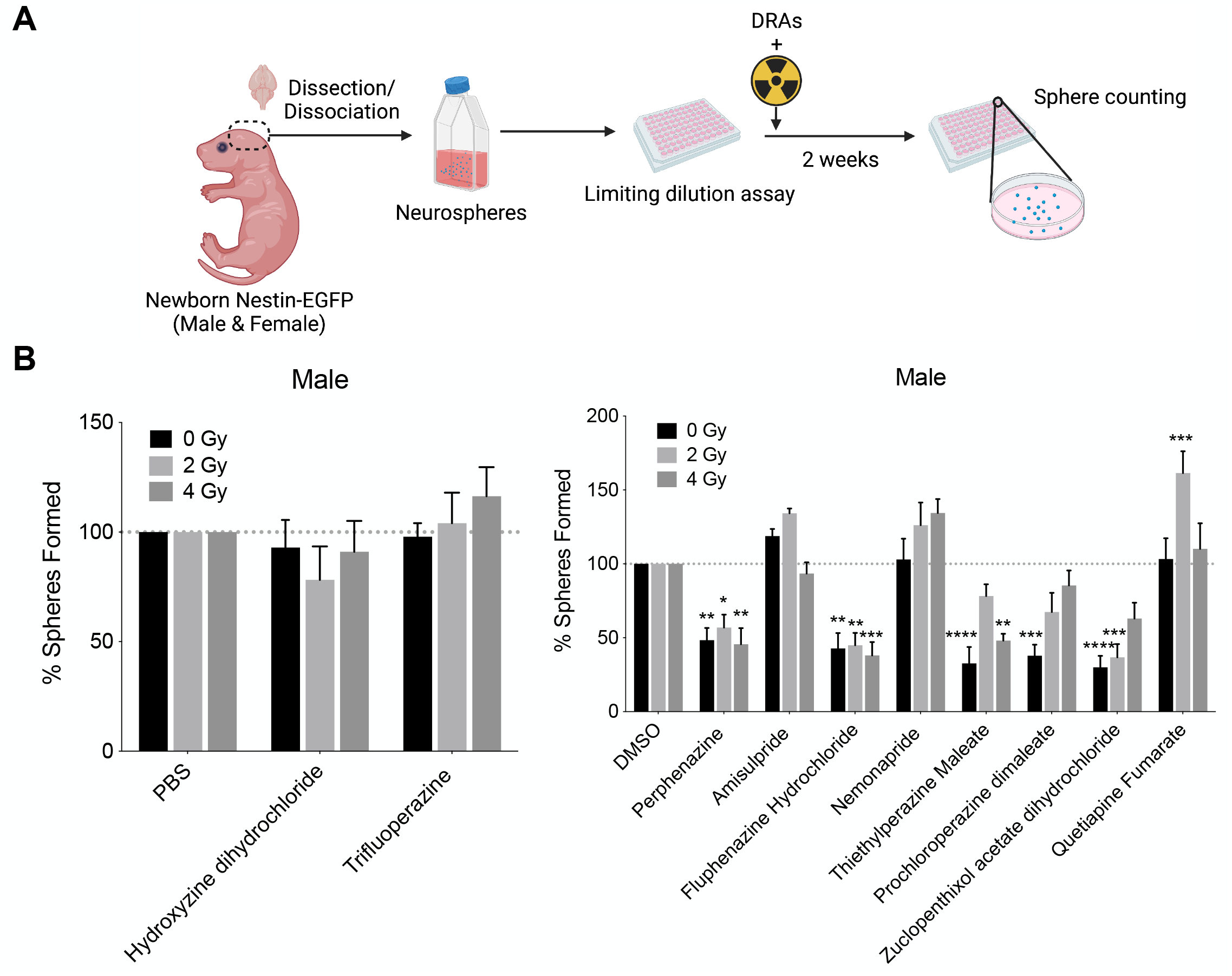
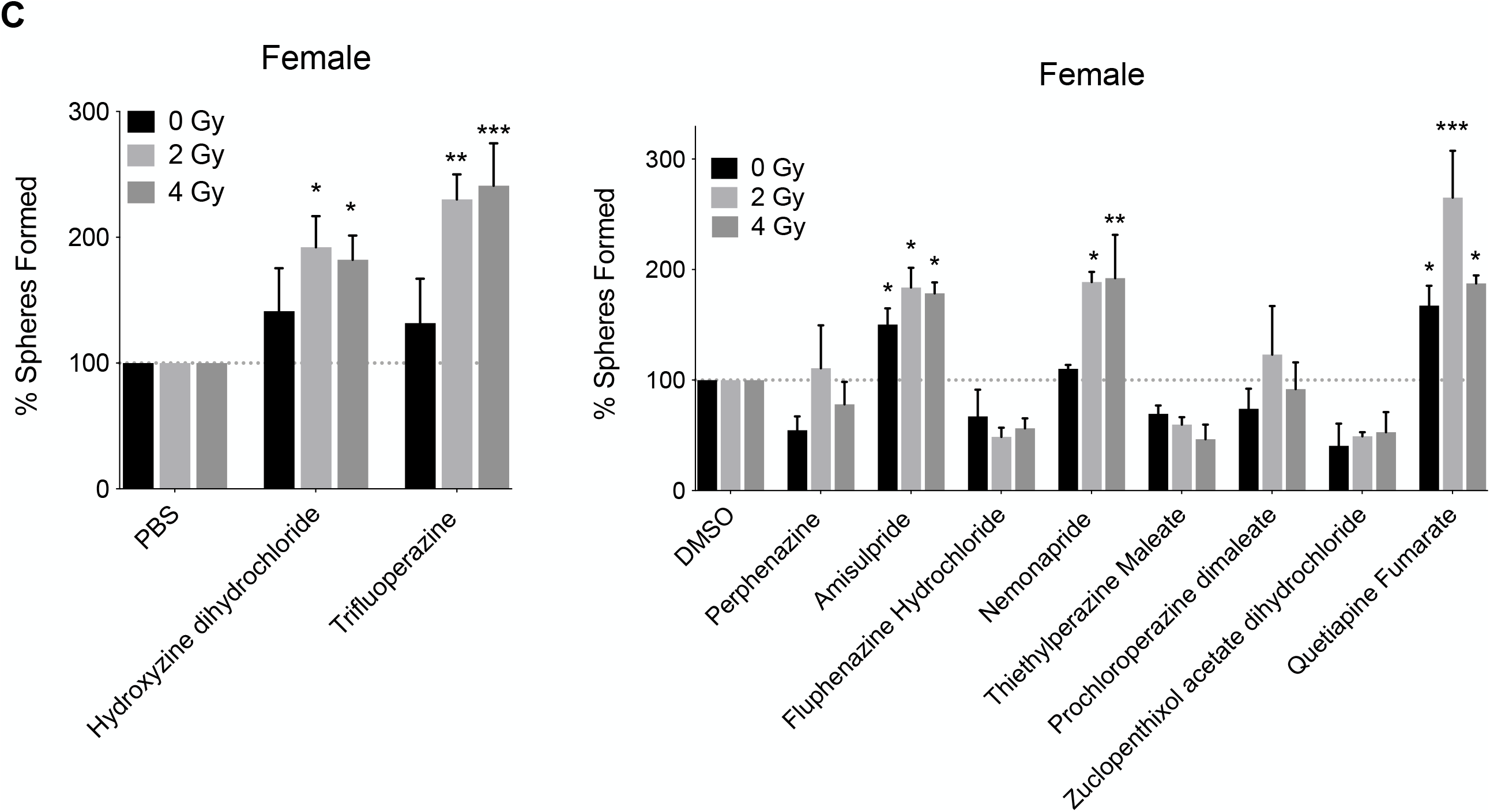
**(A)** Schematic of experimental design underlying Figure 2. **(B, C)** Sphere forming capacity of neurospheres (Passage #2) from male or female newborn pups pre-treated with a single treatment of either controls (DMSO/PBS) or DRAs (1 μM) and subjected to 0, 2 or 4 Gy irradiation. (Unpaired Student’s t-tests. * *p*-value < 0.05, ** *p*-value < 0.01, *** *p*-value < 0.001, **** *p*-value < 0.0001).

Cells from neurospheres established from female mice, treated with hydroxyzine, trifluoperazine, amisulpride, nemonapride, or quetiapine, showed a significant increase in sphere-formation after irradiation with 0, 2, or 4 Gy compared to the corresponding irradiated, solvent-treated controls **(Figure 2C)**.

### Dopamine receptor antagonists show sex-dependent effects on neural stem/progenitor cells in vivo

To test if DRAs that increase neurosphere-formation *in vitro* would also affect neural stem/progenitor cells *in vivo*, we treated 3-week-old Nestin-EGFP mice with five daily i.p. injections of trifluoperazine or solvent and harvested the brains after the last injection **(Figure 3A)**. The brains were digested into single cell suspensions and analyzed for EGFP^+^ NSPCs by flow cytometry. When compared to solvent-treated mice, mice treated with trifluoperazine showed a significant increase in the number of EGFP^+^ NSPCs. The same trend was seen in male mice, but the effect was small and did not reach statistical significance **(Figure 3B)**.

**Figure 3.**
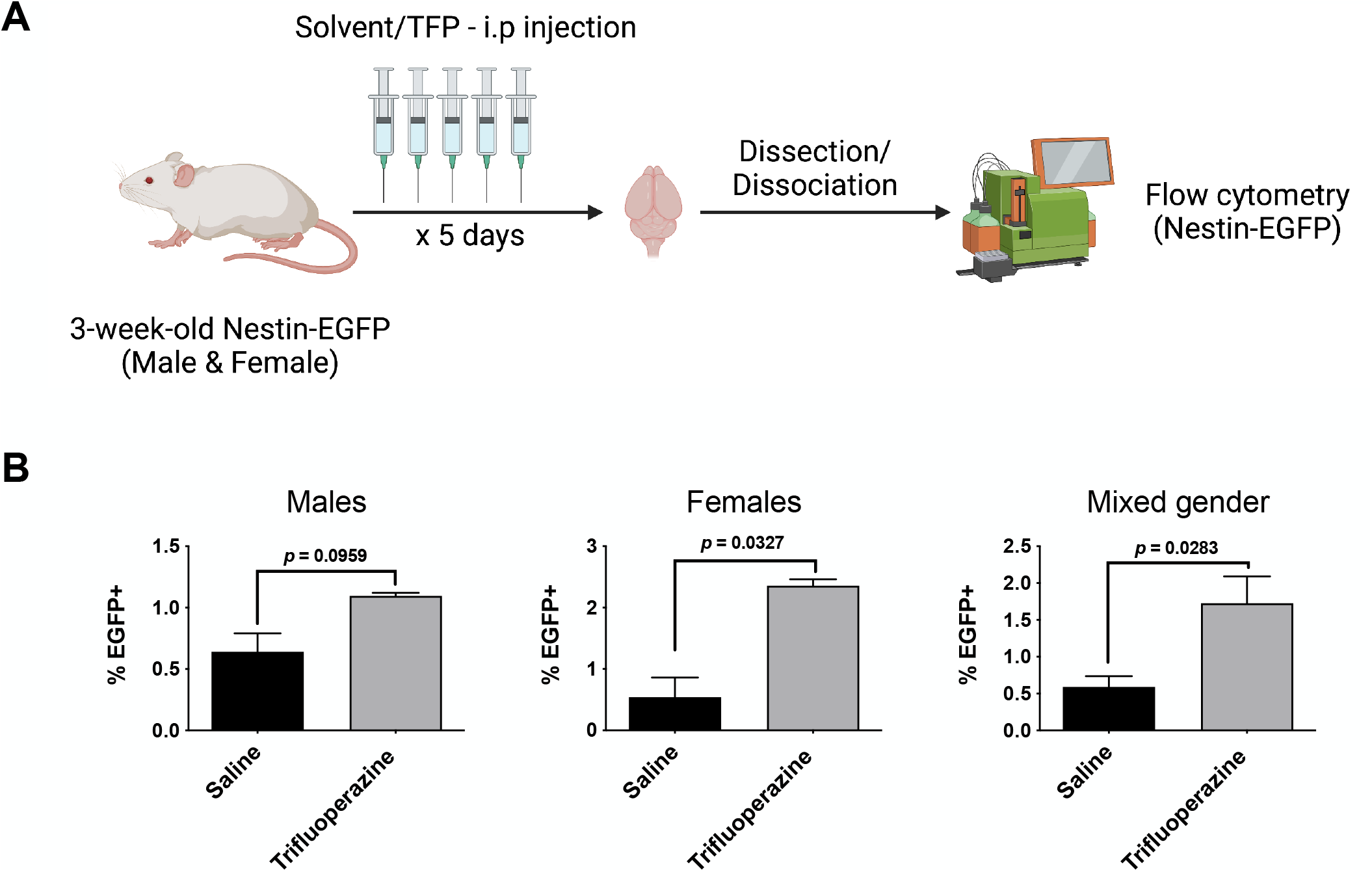
**(A)** Schematic of experimental design underlying Figure 3. **(B)** The percentage of EGFP+ cells in dissociate brain samples from 3-week old Nestin-EGFP male and female mice treated with saline or trifluoperazine (20 mg/kg) i.p. for five days (N=3; Unpaired Student’s t-tests).

Next, we sought to assess if this effect of DRAs could also be observed in 8-week-old mice in which neural stem/progenitor frequencies and the amount of neurogenesis are lower. In 3-week-old mice baseline frequencies of NSPCs in the whole brain were already less than 1 %. Therefore, to enrich frequencies of NSPCs, we collected brain sections from the subventricular zones of the lateral ventricles, regions known to contain stem cells niches, digested the sections into single cells suspensions and analyzed them for the number of Nestin-EGFP^+^ cells by flow cytometry (**Figure 4A**). In cell suspensions from these brain regions we found baseline frequencies of NSPCs reaching 20 % of the total cell population. Treatment with trifluoperazine significantly increased the number of NSPCs in female mice. The same trend was seen in male mice but did not reach the level of statistical significance (**Figure 4B**), which was in line with the findings we reported in 3-week-old mice **(Figure 3B)**. Likewise, treatment of the animals with hydroxyzine, quetiapine, or nemonapride significantly increased the number of NSPCs in combined male and female cohorts, with similar trends seen in the individual sexes (**Figure 4C**). Lastly, treatment with amisulpride significantly increased NSPCs numbers in both sexes (**Figure 4D, 5A/B).**

**Figure 4.**
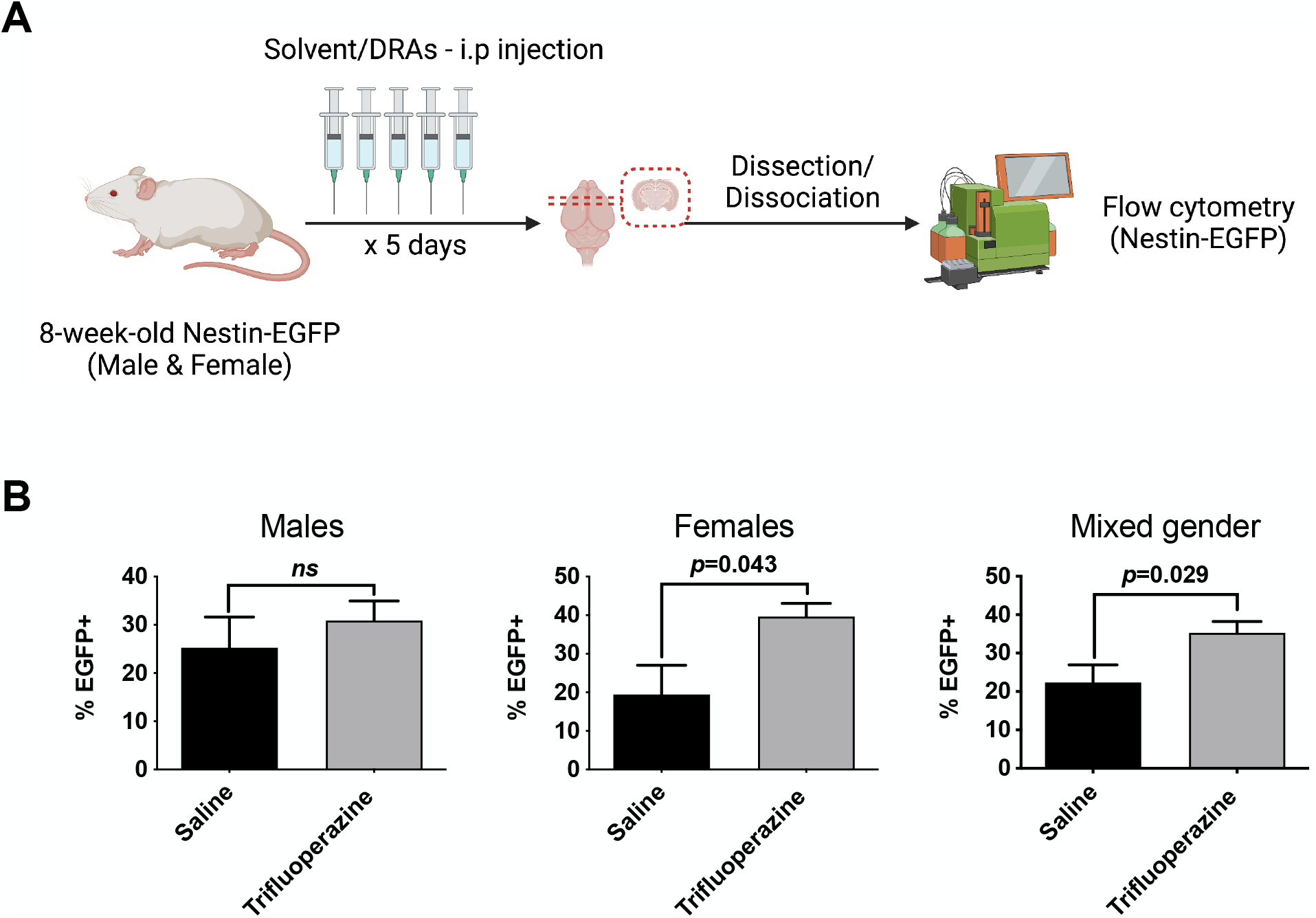
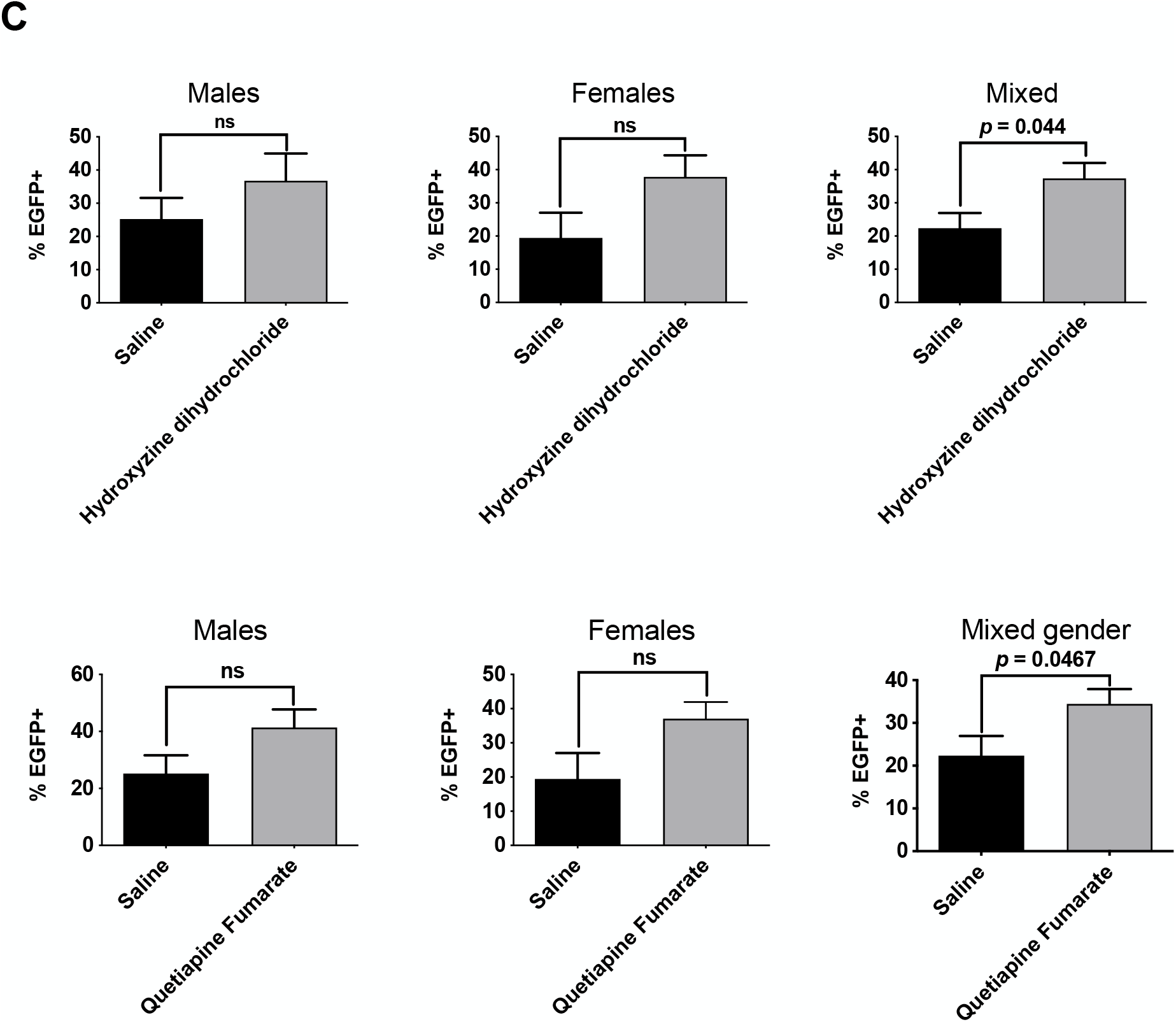
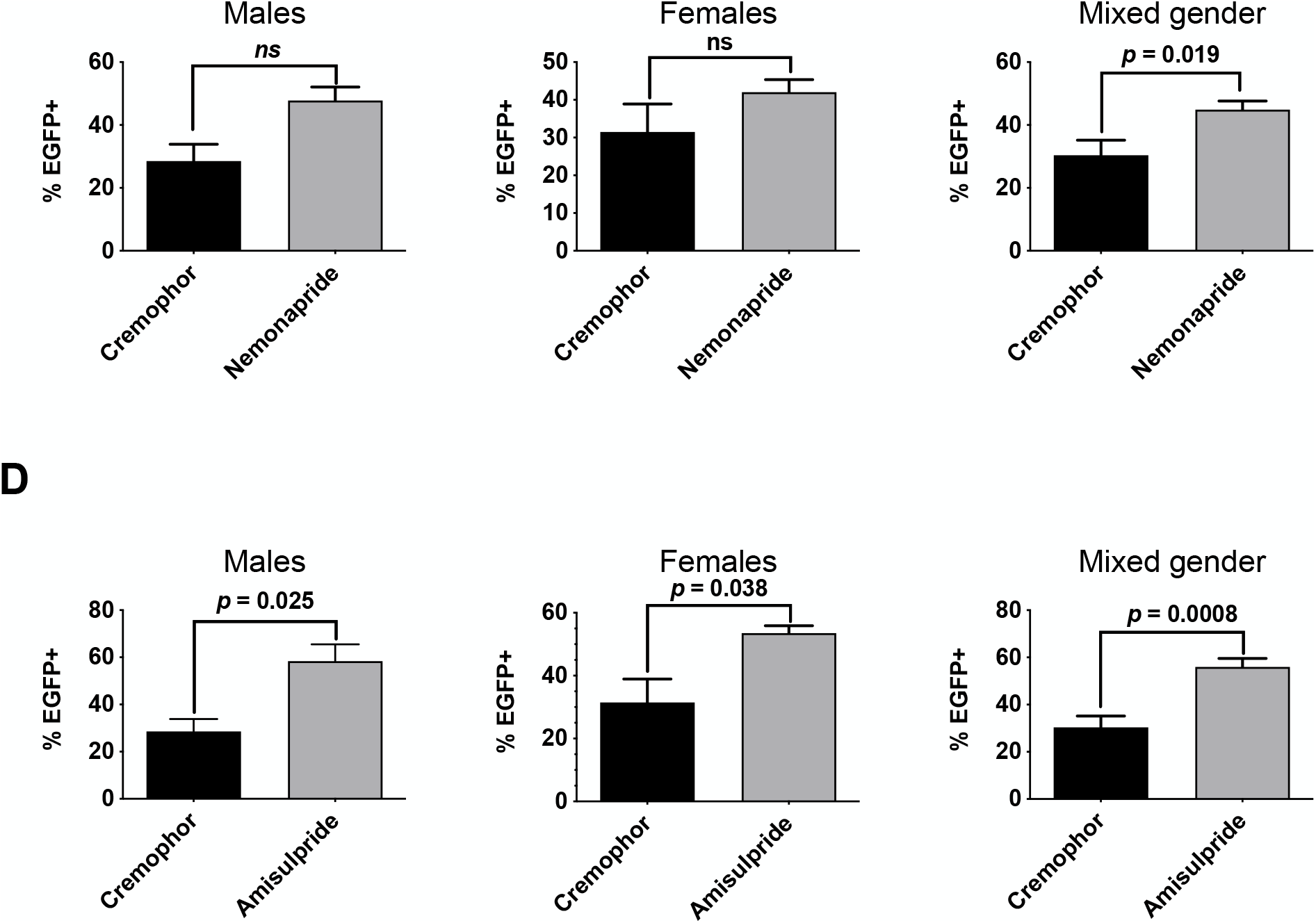
**(A)** Schematic of experimental design underlying Figure 4 **(B-D)** The percentage of EGFP+ cells in dissociate brain samples from 8-week-old Nestin-EGFP male and female mice treated with either Control (Saline/Cremophor) or DRAs (20 mg/kg of trifluoperazine, 30 mg/kg of quetiapine, 20 mg/kg of Hydroxyzine dihydrochloride, 1 mg/kg of Nemonapride or 1 mg/kg of Amisulpride) i.p. for five days. (N=3; Unpaired Student’s t-tests).

**Figure 5.**
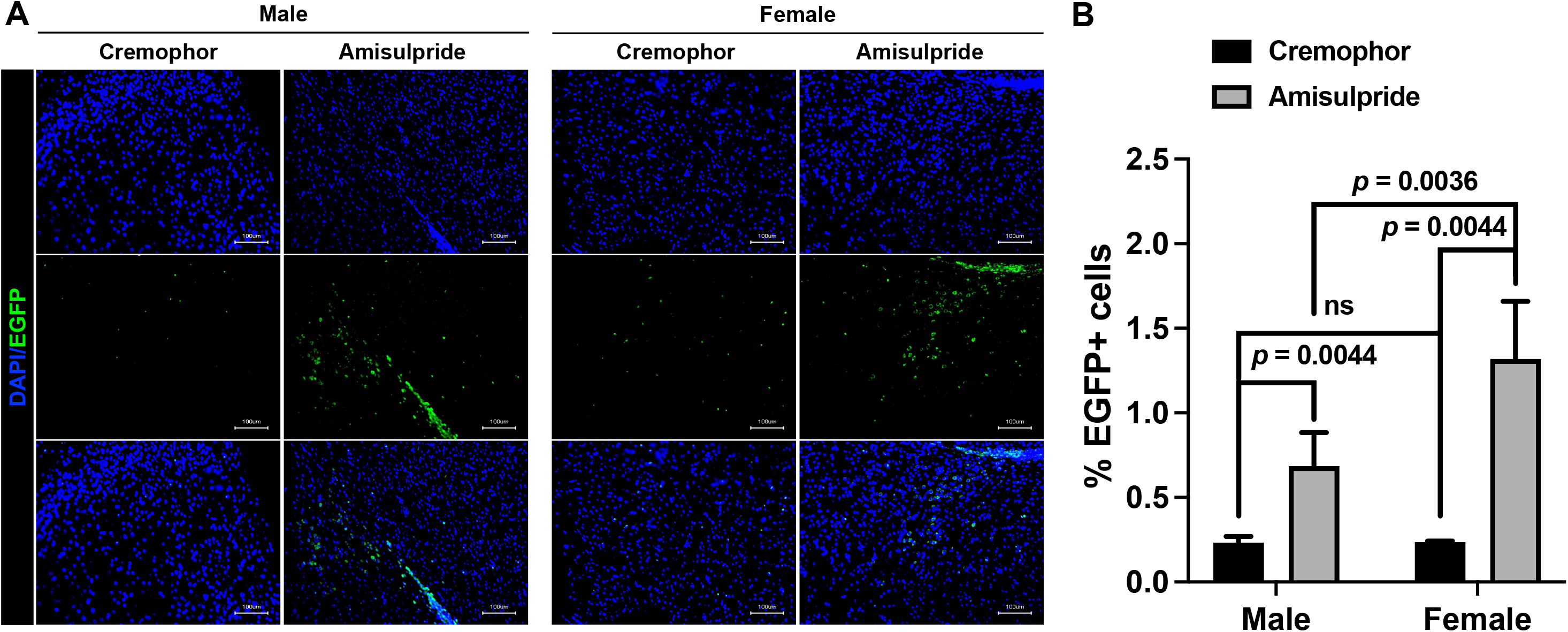
**(A)** Immunofluorescent staining for EGFP in coronal sections of brains from 8-week-old Nestin-EGFP male and female mice treated with either Cremophor (control) or 1 mg/kg of Amisulpride i.p. for five days. **(B)** Quantification of the percentage of EGFP+ cells over DAPI-stained cells. (Unpaired Student’s t-tests).

## Discussion

Glioblastoma (GBM) is a deadly form of brain cancer. The current standard of care, surgery followed by radiotherapy and chemotherapy only provides a limited advantage to patients, with the 5-year survival rate for pediatric patients being 20 % and less than 5 % for adults. These unacceptably low rates highlight urgent need for novel treatment strategies. So far, targeted agents and biologicals have not been successful in meeting this need, often due to their inability to cross the blood-brain barrier (BBB). In a high-throughput screen (8) we previously identified dopamine-receptor antagonists as FDA-approved drugs, BBB-penetrating drugs able to target glioma-initiating cells in mouse models of GBM and lead to significantly improved median overall survival (6, 8, 9). The atypical dopamine receptor antagonist and ClpP activator ONC201 has entered clinical trials against GBM (16, 17), including for pediatric patients suffering from H3 K27M-mutant diffuse midline glioma (7). A monotherapy with dopamine receptor antagonist has little to no effect on GBM and therefore, dopamine receptor antagonists are likely combined with radiotherapy or chemotherapy. Radiotherapy and chemotherapy both affect cognitive function especially in pediatric patients (18–20). Therefore, we sought to test if dopamine receptor antagonists affect normal stem cells in the CNS alone or in combination with radiation.

Our data indicate a trend for higher baseline numbers of NSPC in female newborn mice when compared to age-matched male mice. Even though the variability between batches of mice was large and the differences did not reach statistical significance, the observed sex differences were consistent. This agreed with previous studies that explored the role of sexual dimorphism in regulating the ventricular-subventricular zone (V-SVZ), one of the locations in which neural stem cells (NSCs) reside (21, 22). Neurogenesis was found to be more prominent in female mice compared to age-matched males with higher proliferating rates and lower numbers of apoptotic cells seen in the V-SVZ of adult mice (22). Moreover, males showed significant decline in NSPC numbers, progenitor cell proliferation, and more disorganized migrating neuroblast chains with age when compared to age-matched females, indicating sex differences indeed exist in the process of neurogenesis and decline with age, at least in adults (21).

Dopamine receptor antagonists belong to a broad class of psychotropic drugs with varying specificity for dopamine, serotonin, alpha- and beta-adrenergic, histamine, and muscarinic receptors, thus leading to different spectra of target and off-target effects. We previously observed a broad range of efficacies for different dopamine receptor antagonists in preventing or promoting cancer cell plasticity characterized by the acquisition of a GIC phenotype in surviving, previously non-stem glioma cells in response to radiation (6, 8, 9). E.g., quetiapine and trifluoperazine prevented radiationinduction of cancer-initiating cells from non-stem glioma cells, while amisulpride promoted the plasticity of non-stem cancer cells.

While our screen was performed with glioblastoma cells of male origin, but quetiapine and trifluoperazine also showed efficacy in glioblastoma and breast cancer cells obtained from female patients. In the present study, we expanded our study taking sex differences in normal stem cells into consideration. Hydroxyzine, trifluoperazine, amisulpride, nemonapride or quetiapine alone or in combination with radiation significantly increased the number of NSPCs in Passage #2 neurospheres from female but not from male mice.

This was in stark contrast to our studies using the dopamine receptor antagonists trifluoperazine and quetiapine in glioma where we found both drugs preventing glioma cell plasticity and targeting glioblastoma stem cells (GSCs) (8, 9). The differential response of GSCs and NSPCs is most likely grounded in their differential reliance on developmental signaling pathways. In our previous studies, GSCs relied on Yamanaka factors (Sox2, Klf4, c-Myc, and Oct4), which were induced by radiation. This induction was prevented by trifluoperazine or quetiapine in a DRD2-dependent manner (8, 9), which led to a loss of the inactivating Ser9/21 phosphorylation of Glycogen synthase kinase 3α/β (GSK3α/β), subsequent degradation of β-catenin, reduced self-renewal capacity of patient-derived GBM lines and prolonged median survival in patient-derived orthotopic xenograft (PDOX) bearing mice (8). While dependent on Wnt/β-catenin signaling (23–25), NSCs are maintained by additional developmental signaling pathways, including Notch (26–28) and Shh (29, 30), and these pathways are also modulated by GSK3α/β (31, 32).

In the brain, both GSK3α and GSK3β are expressed (33), and disruption of GSK3 signaling has been implicated in a number of neurological diseases, such as schizophrenia (34), bipolar disorder (35), and neurodegenerative diseases (36). However, studies have shown that GSK3α and GSK3β play overlapping but distinct roles in neocortex development (37), with GSK3β acting as the master kinase in the regulation of neural stem/progenitor cell behavior (38). The GSK3β-dependent regulation of stem cells includes the maintenance of self-renewal (39), and the control of cell differentiation (40, 41), and the plasticity for stem cell survival (42), all critical characteristics for stem cell survival. To be noted, certain types of stem cells require a very tight control of GSK3β activity, since aberrant expression can lead to very distinct outcomes, such as is the case with neural stem cells being pushed to cell death by means of autophagy following over-activation of GSK3β (43). Also, GSK3α/β signaling is known to have differential effects in naïve and primed embryonic stem cells (44, 45). Our previous studies suggested that radiation induced a transient multipotent state in surviving GBM cells through epigenetic remodeling (8, 46). This induced multipotent state could explain the differential effects of dopamine receptor antagonists on GBM cells and lineage committed NSCs, given the potential differences in the GSK3α/β signaling for those two cell populations.

A limitation of our study is its restriction to a murine *in vivo* model. Species differences in the dependence of corresponding stem cell populations on developmental pathways have been reported before (44, 47, 48). While the effects of some dopamine receptor antagonists and selected other psychotropic drugs have been tested in human NSPCs, *in vivo* responses will most likely also include effects on the microenvironment and cannot easily be assessed in the human setting.

To the best of our knowledge, our study is the first to investigate the combined effects of dopamine receptor antagonists and radiation in NSPCs. The unexpected finding that dopamine receptor antagonists either protected irradiated NSPCs from radiation or caused an expansion of the NSPC pool indicates that this effective combination therapy against GBM (6, 8, 9) will not increase radiation-induced cognitive decline through further increasing the elimination of NSPCs and subsequent loss of neurogenesis.

In conclusion, our published data (8, 9) and data presented in this study suggest the existence of a therapeutic window for dopamine receptor antagonists in combination with radiation as a novel combination therapy against GBM. Normal tissue toxicity of this combination potentially differs depending on age and sex and should be taken into consideration when designing clinical trials.

